## Supplementary Information for "Polygenic Prediction via Bayesian Regression and Continuous Shrinkage Priors"

[1]Psychiatric and Neurodevelopmental Genetics Unit, Center for Genomic Medicine, Massachusetts General Hospital, Boston, MA 02114, USA; [2]Department of Psychiatry, Massachusetts General Hospital, Harvard Medical School, Boston, MA 02114, USA; [3]Stanley Center for Psychiatric Research, Broad Institute of MIT and Harvard, Cambridge, MA 02138, USA; [4]Analytic and Translational Genetics Unit, Center for Genomic Medicine, Massachusetts General Hospital, Boston, MA 02114, USA; [5]Department of Statistics, Texas A&M University, College Station, TX 77843, USA

Address correspondence to:

Tian Ge

Psychiatric and Neurodevelopmental Genetics Unit

Center for Genomic Medicine

Massachusetts General Hospital

#### Supplementary Note

The Bayesian regression model for PRS-CS and PRS-CS-auto is:

$$\begin{aligned} &\boldsymbol{y} = \boldsymbol{Z}\boldsymbol{\beta} + \boldsymbol{\epsilon}, \qquad \boldsymbol{\epsilon} \sim \mathrm{N}(\mathbf{0}, \sigma^2 \boldsymbol{I}), \qquad p(\sigma^2) \propto \sigma^{-2}, \\ &\beta_j \sim \mathrm{N}\left(0, \frac{\sigma^2}{N}\psi_j\right), \qquad \psi_j \sim \mathrm{G}(a, \delta_j), \qquad \delta_j \sim \mathrm{G}(b, \phi), \end{aligned} \tag{1}$$

where $\boldsymbol{y}$ and $\boldsymbol{Z}$ have been standardized. The full conditional distributions for all the parameters in this model are analytically tractable, and thus an efficient Gibbs sampler can be derived.

Let $\mathrm{MVN}(\boldsymbol{\mu}, \boldsymbol{\Sigma})$ denote the multivariate normal distribution with mean $\boldsymbol{\mu}$ and covariance $\boldsymbol{\Sigma}$; $\mathrm{G}(\alpha, \beta)$ and $\mathrm{iG}(\alpha, \beta)$ denote the gamma distribution and inverse-gamma distribution with shape parameter $\alpha$ and scale parameter $\beta$, respectively; and $\mathrm{giG}(p, \rho, \chi)$ denote the three-parameter generalized inverse Gaussian distribution with probability density function

$$f(x; \lambda, \rho, \chi) = \frac{(\rho/\chi)^{\lambda/2}}{2K_\lambda(\sqrt{\rho\chi})} x^{\lambda-1} \mathrm{e}^{-(\rho x + \chi/x)/2}, \qquad x > 0, \quad \rho > 0, \quad \chi > 0, \tag{2}$$

where $K_\lambda$ is the modified Bessel function of the second kind. Let $N$ and $M$ denote the sample size and the total number of genetic markers, respectively. In addition, let $\widehat{\boldsymbol{\beta}} = \boldsymbol{Z}^\top \boldsymbol{y}/N$ denote the marginal least squares effect size estimates from the genome-wide association study, $\boldsymbol{\Psi} = \mathrm{diag}\{\psi_1, \psi_2, \cdots, \psi_M\}$, and $\boldsymbol{D} = \boldsymbol{Z}^\top \boldsymbol{Z}/N$ denote the LD matrix. The Gibbs sampler then involves the following steps in each MCMC iteration:

- update $\boldsymbol{\beta}$: $\quad [\, \boldsymbol{\beta} \mid \sigma^2, \boldsymbol{\Psi}, \widehat{\boldsymbol{\beta}}, \boldsymbol{D} \,] \sim \mathrm{MVN}(\boldsymbol{\mu}, \boldsymbol{\Sigma}), \quad \boldsymbol{\mu} = \frac{N}{\sigma^2}\boldsymbol{\Sigma}\widehat{\boldsymbol{\beta}}, \quad \boldsymbol{\Sigma} = \frac{\sigma^2}{N}(\boldsymbol{D} + \boldsymbol{\Psi}^{-1})^{-1},$
- update $\sigma^2$: $\quad [\, \sigma^2 \mid \boldsymbol{\beta}, \boldsymbol{\Psi}, \widehat{\boldsymbol{\beta}}, \boldsymbol{D} \,] \sim \mathrm{iG}\left(\frac{N+M}{2}, \frac{N}{2}\left[1 - 2\boldsymbol{\beta}^\top \widehat{\boldsymbol{\beta}} + \boldsymbol{\beta}^\top (\boldsymbol{D} + \boldsymbol{\Psi}^{-1})\boldsymbol{\beta}\right]\right),$
- update $\psi_j$: $\quad [\, \psi_j \mid \beta_j, \sigma^2, \delta_j \,] \sim \mathrm{giG}\left(a - \frac{1}{2}, 2\delta_j, \frac{N\beta_j^2}{\sigma^2}\right),$
- update $\delta_j$: $\quad [\, \delta_j \mid \psi_j \,] \sim \mathrm{G}(a + b, \psi_j + \phi).$

In PRS-CS-auto, we assign a half-Cauchy prior on the global shrinkage parameter[1,2]: $\phi^{1/2} \sim \mathrm{C}^+(0, 1)$, which is equivalent to a scale mixture of gamma distributions: $\phi \sim \mathrm{G}(1/2, \omega)$, $\omega \sim \mathrm{G}(1/2, 1)$. Gibbs updates can then be derived using this augmented representation:

- update $\phi$: $\quad [\, \phi \mid \delta_j, \omega \,] \sim \mathrm{G}\left(Mb + \frac{1}{2}, \sum_{j=1}^{M} \delta_j + \omega\right),$
- update $\omega$: $\quad [\, \omega \mid \phi \,] \sim \mathrm{G}(1, 1 + \phi).$

We generate random variates from the generalized inverse Gaussian distribution using the algorithm described in Devroye[3]. We note that $\boldsymbol{y}$ and $\boldsymbol{Z}$ did not appear in any of the updates above, and thus individual-level data is not required for model fitting. In practice, $\boldsymbol{D}$ and $\boldsymbol{\Psi}$ are $M \times M$ matrices, and the calculation of $(\boldsymbol{D} + \boldsymbol{\Psi}^{-1})^{-1}$ becomes computationally infeasible when $M$ is large. We thus partition the genome into 1,703 largely independent genomic regions estimated using data from the 1000 Genomes Project European sample[4], and in each MCMC iteration sequentially update the SNP effect sizes within each LD block $\ell$:

$$[\, \boldsymbol{\beta}_\ell \mid \sigma^2, \boldsymbol{\Psi}_\ell, \widehat{\boldsymbol{\beta}}_\ell, \boldsymbol{D}_\ell \,] \sim \mathrm{MVN}(\boldsymbol{\mu}_\ell, \boldsymbol{\Sigma}_\ell), \quad \boldsymbol{\mu}_\ell = \frac{N}{\sigma^2} \boldsymbol{\Sigma}_\ell \widehat{\boldsymbol{\beta}}_\ell, \quad \boldsymbol{\Sigma}_\ell = \frac{\sigma^2}{N} (\boldsymbol{D}_\ell + \boldsymbol{\Psi}_\ell^{-1})^{-1}. \tag{3}$$

The LD matrix $\boldsymbol{D}_\ell$ for each LD block can be estimated using an external reference panel.

### Supplementary Figures

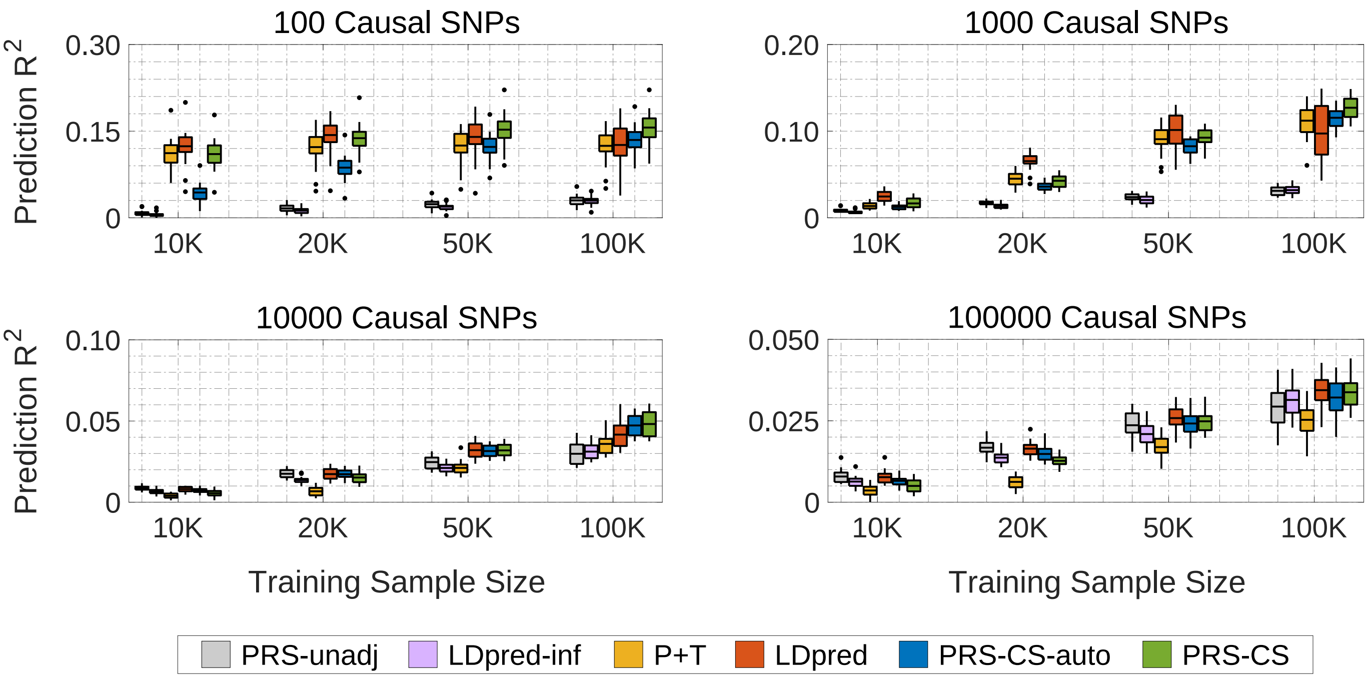


**Supplementary Figure 1: Predictive performance of six polygenic prediction methods in simulation studies using a point-normal model with heritability fixed at 0.2**. The 1000 Genomes Project European sample was used as an external linkage disequilibrium (LD) reference panel. Tuning parameters ($P$-value threshold in P+T, fraction of causal SNPs in LDpred, and global shrinkage parameter in PRS-CS) were selected in a validation data set. Prediction accuracy was quantified by $R^2$ between the observed and predicted traits in an independent testing set. The four panels correspond to the four genetic architectures (100, 1,000, 10,000 and 100,000 causal variants) simulated using the point-normal model. Within each panel, results for four different training sample sizes (10,000, 20,000, 50,000 and 100,000) are shown. On each box, the central mark is the mean across 20 simulations, the edges of the box are the 25th and 75th percentiles, the whiskers extend to the most extreme data points that are not considered outliers, and the outliers are plotted individually.

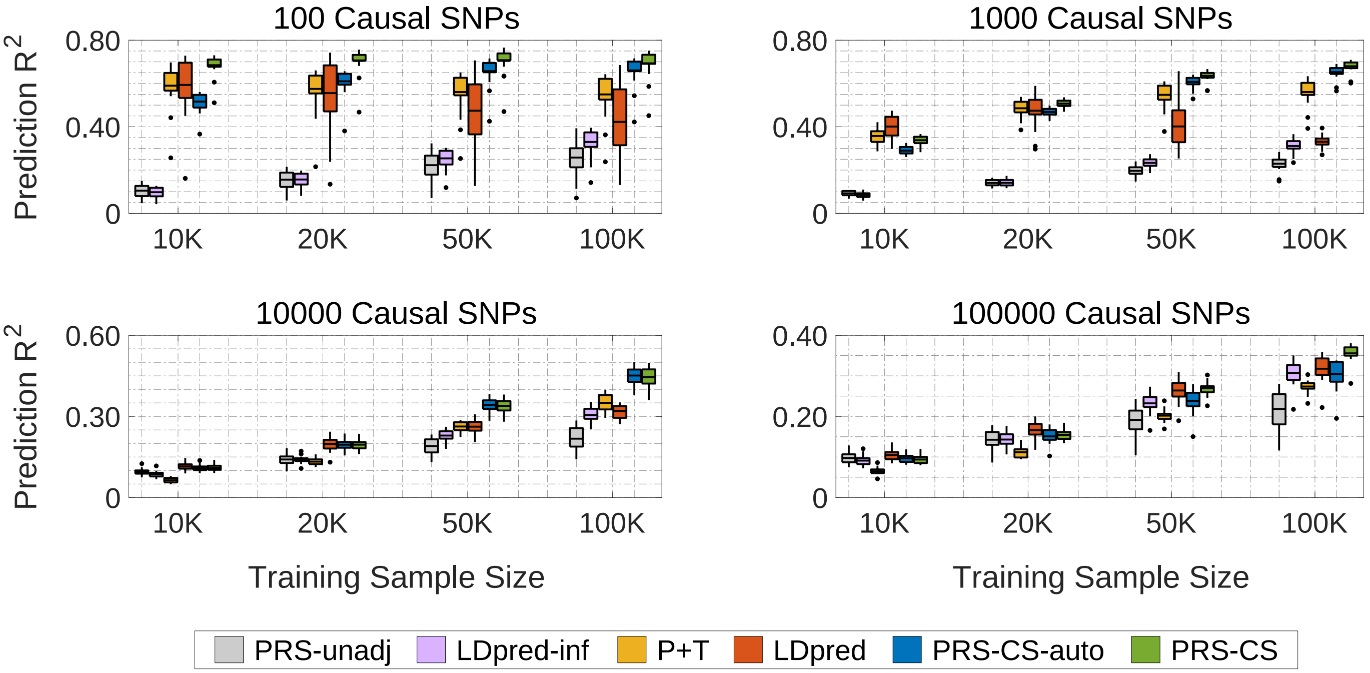


**Supplementary Figure 2: Predictive performance of six polygenic prediction methods in simulation studies using a point-normal model with heritability fixed at 0.8**. The 1000 Genomes Project European sample was used as an external linkage disequilibrium (LD) reference panel. Tuning parameters ($P$-value threshold in P+T, fraction of causal SNPs in LDpred, and global shrinkage parameter in PRS-CS) were selected in a validation data set. Prediction accuracy was quantified by $R^2$ between the observed and predicted traits in an independent testing set. The four panels correspond to the four genetic architectures (100, 1,000, 10,000 and 100,000 causal variants) simulated using the point-normal model. Within each panel, results for four different training sample sizes (10,000, 20,000, 50,000 and 100,000) are shown. On each box, the central mark is the mean across 20 simulations, the edges of the box are the 25th and 75th percentiles, the whiskers extend to the most extreme data points that are not considered outliers, and the outliers are plotted individually.

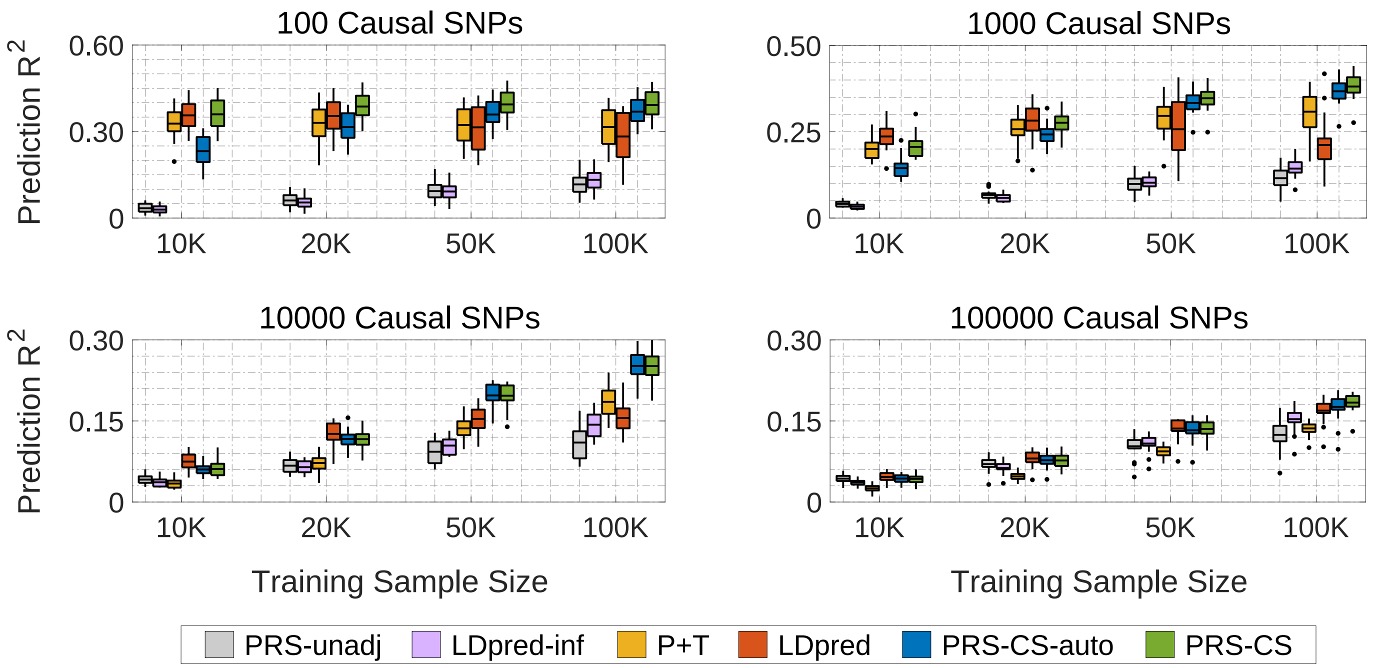


**Supplementary Figure 3: Predictive performance of six polygenic prediction methods in simulation studies using a point-$t$ model with heritability fixed at 0.5**. The 1000 Genomes Project European sample was used as an external linkage disequilibrium (LD) reference panel. Tuning parameters ($P$-value threshold in P+T, fraction of causal SNPs in LDpred, and global shrinkage parameter in PRS-CS) were selected in a validation data set. Prediction accuracy was quantified by $R^2$ between the observed and predicted traits in an independent testing set. The four panels correspond to the four genetic architectures (100, 1,000, 10,000 and 100,000 causal variants) simulated using the point-$t$ model (a mixture of a point mass at zero and a Student's $t$-distribution with 4 degrees of freedom). Within each panel, results for four different training sample sizes (10,000, 20,000, 50,000 and 100,000) are shown. On each box, the central mark is the mean across 20 simulations, the edges of the box are the 25th and 75th percentiles, the whiskers extend to the most extreme data points that are not considered outliers, and the outliers are plotted individually.

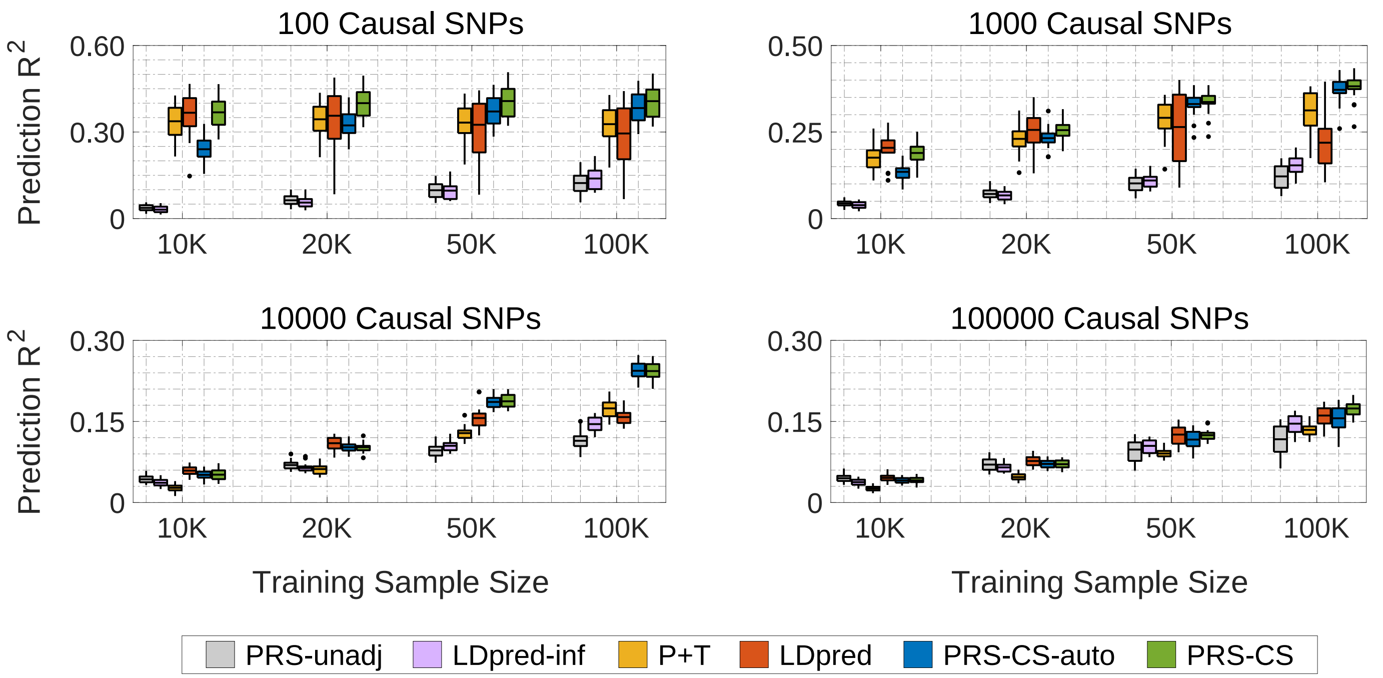


**Supplementary Figure 4: Predictive performance of six polygenic prediction methods in simulation studies using a point-gamma model with heritability fixed at 0.5**. The 1000 Genomes Project European sample was used as an external linkage disequilibrium (LD) reference panel. Tuning parameters ($P$-value threshold in P+T, fraction of causal SNPs in LDpred, and global shrinkage parameter in PRS-CS) were selected in a validation data set. Prediction accuracy was quantified by $R^2$ between the observed and predicted traits in an independent testing set. The four panels correspond to the four genetic architectures (100, 1,000, 10,000 and 100,000 causal variants) simulated using the point-gamma model (a mixture of a point mass at zero and a gamma distribution with the shape parameter set to 2). Within each panel, results for four different training sample sizes (10,000, 20,000, 50,000 and 100,000) are shown. On each box, the central mark is the mean across 20 simulations, the edges of the box are the 25th and 75th percentiles, the whiskers extend to the most extreme data points that are not considered outliers, and the outliers are plotted individually.

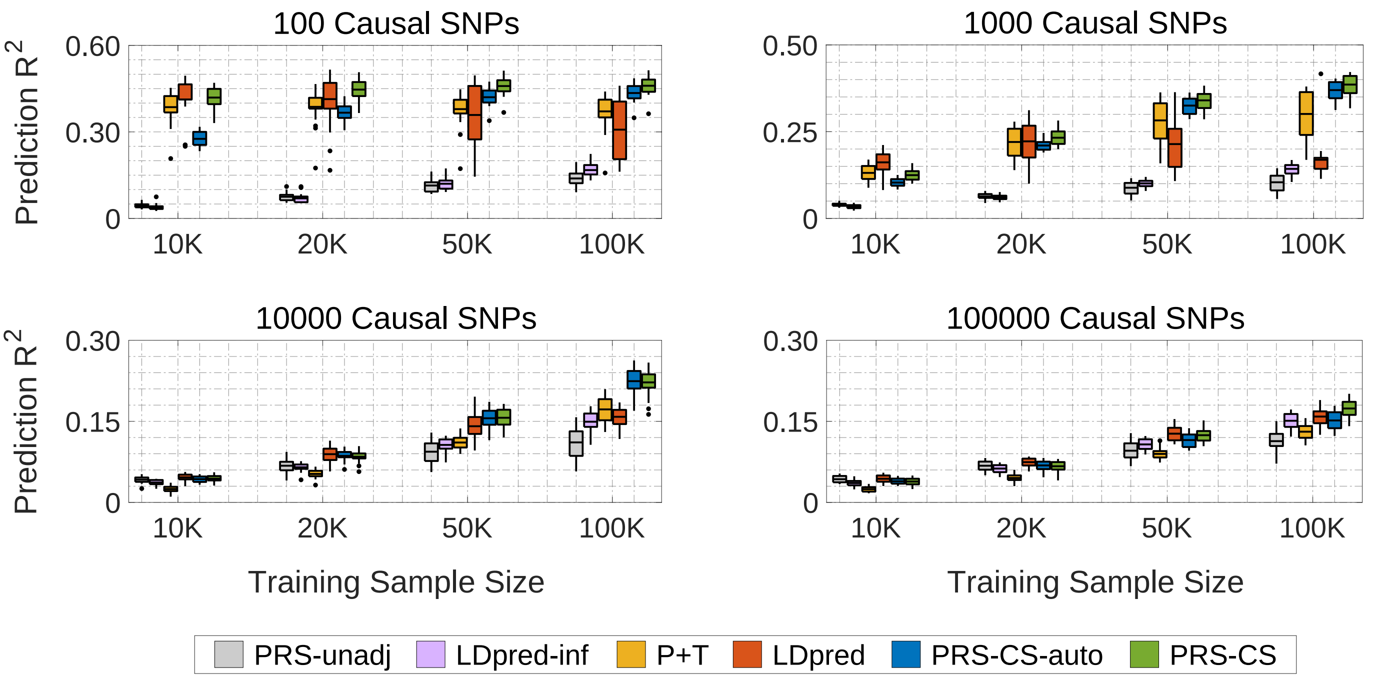


**Supplementary Figure 5: Predictive performance of six polygenic prediction methods in simulation studies using an in-sample reference panel**. SNP effect sizes were simulated using a point-normal model with different numbers of causal variants and heritability was fixed at 0.5. The combined validation and testing data sets ($N = 6,000$) were used as an in-sample linkage disequilibrium (LD) reference panel. Tuning parameters ($P$-value threshold in P+T, fraction of causal SNPs in LDpred, and global shrinkage parameter in PRS-CS) were selected in a validation data set. Prediction accuracy was quantified by $R^2$ between the observed and predicted traits in an independent testing set. The four panels correspond to the four genetic architectures (100, 1,000, 10,000 and 100,000 causal variants) simulated using the point-normal model. Within each panel, results for four different training sample sizes (10,000, 20,000, 50,000 and 100,000) are shown. On each box, the central mark is the mean across 20 simulations, the edges of the box are the 25th and 75th percentiles, the whiskers extend to the most extreme data points that are not considered outliers, and the outliers are plotted individually.

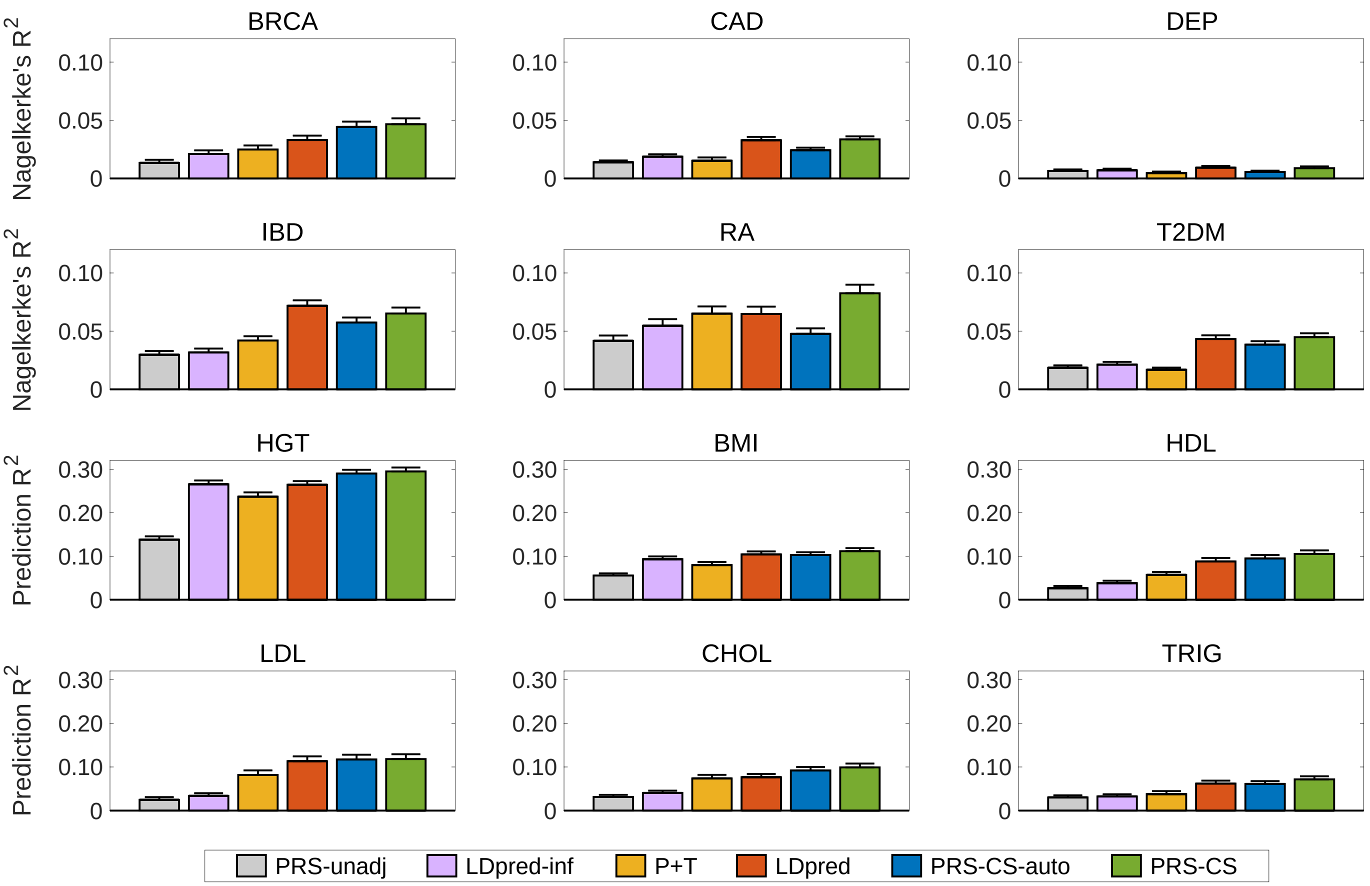


**Supplementary Figure 6: Prediction accuracy of six polygenic prediction methods in the Partners HealthCare Biobank using an in-sample reference panel**. Posterior SNP effect sizes were trained with large-scale genome-wide association summary statistics, using the Partners HealthCare Biobank data ($N = 19,136$) as an in-sample linkage disequilibrium (LD) reference panel. Polygenic scores were applied to predict six curated common complex diseases — breast cancer (BRCA), coronary artery disease (CAD), depression (DEP), inflammatory bowel disease (IBD), rheumatoid arthritis (RA), and type 2 diabetes mellitus (T2DM), and six quantitative traits — height (HGT), body mass index (BMI), high-density lipoproteins (HDL), low-density lipoproteins (LDL), cholesterol (CHOL), and triglycerides (TRIG). The Partners HealthCare Biobank sample for each disease and quantitative phenotype was repeatedly and randomly split into a validation set comprising 1/3 of the data and a testing set comprising 2/3 of the data. Tuning parameters ($P$-value threshold in P+T, fraction of causal SNPs in LDpred, and global shrinkage parameter in PRS-CS) were selected in the validation data set, and the predictive performance was assessed in the testing set. For disease (case-control) phenotypes and quantitive traits, prediction accuracy was measured by the Nagelkerke's $R^2$ and $R^2$, respectively, averaged across 100 random splits. The error bar indicates the standard deviation of prediction accuracy across 100 random splits.

#### Supplementary Tables

**Supplementary Table 1:** Numerical results of the simulation studies shown in Fig. 1. SNP effect sizes were simulated using a point-normal model with different numbers of causal variants or a normal mixture model. Heritability was fixed at 0.5. The 1000 Genomes Project European sample was used as an external linkage disequilibrium (LD) reference panel. For each combination of the genetic architecture and the training sample size, the mean and standard deviation of the prediction accuracy for each polygenic prediction method in the testing data set across 20 simulations are reported.

**Supplementary Table 2:** Numerical results of the simulation studies shown in Supplementary Fig. 1. SNP effect sizes were simulated using a point-normal model with different numbers of causal variants. Heritability was fixed at 0.2. The 1000 Genomes Project European sample was used as an external linkage disequilibrium (LD) reference panel. For each combination of the genetic architecture and the training sample size, the mean and standard deviation of the prediction accuracy for each polygenic prediction method in the testing data set across 20 simulations are reported.

**Supplementary Table 3:** Numerical results of the simulation studies shown in Supplementary Fig. 2. SNP effect sizes were simulated using a point-normal model with different numbers of causal variants. Heritability was fixed at 0.8. The 1000 Genomes Project European sample was used as an external linkage disequilibrium (LD) reference panel. For each combination of the genetic architecture and the training sample size, the mean and standard deviation of the prediction accuracy for each polygenic prediction method in the testing data set across 20 simulations are reported.

**Supplementary Table 4:** Numerical results of the simulation studies shown in Supplementary Fig. 3. SNP effect sizes were simulated using a point-$t$ model (a mixture of a point mass at zero and a Student's $t$-distribution with 4 degrees of freedom) with different numbers of causal variants. Heritability was fixed at 0.5. The 1000 Genomes Project European sample was used as an external linkage disequilibrium (LD) reference panel. For each combination of the genetic architecture and the training sample size, the mean and standard deviation of the prediction accuracy for each polygenic prediction method in the testing data set across 20 simulations are reported.

**Supplementary Table 5:** Numerical results of the simulation studies shown in Supplementary Fig. 4. SNP effect sizes were simulated using a point-gamma model (a mixture of a point mass at zero and a gamma distribution with the shape parameter set to 2) with different numbers of causal variants. Heritability was fixed at 0.5. The 1000 Genomes Project European sample was used as an external linkage disequilibrium (LD) reference panel. For each combination of the genetic architecture and the training sample size, the mean and standard deviation of the prediction accuracy for each polygenic prediction method in the testing data set across 20 simulations are reported.

**Supplementary Table 6:** Numerical results of the simulation studies shown in Supplementary Fig. 5. SNP effect sizes were simulated using a point-normal model with different numbers of causal variants. Heritability was fixed at 0.5. The combined validation and testing data sets were used as an in-sample linkage disequilibrium (LD) reference panel. For each combination of the genetic architecture and the training sample size, the mean and standard deviation of the prediction accuracy for each polygenic prediction method in the testing data set across 20 simulations are reported.

**Supplementary Table 7:** Calibration slopes (the slope of regressing the true phenotype onto the polygenic score predictor) for the six polygenic prediction methods evaluated in Fig. 1 and Supplementary Table 1.

**Supplementary Table 8:** Calibration slopes (the slope of regressing the true phenotype onto the polygenic score predictor) for the six polygenic prediction methods evaluated in Supplementary Fig. 1 and Supplementary Table 2.

**Supplementary Table 9:** Calibration slopes (the slope of regressing the true phenotype onto the polygenic score predictor) for the six polygenic prediction methods evaluated in Supplementary Fig. 2 and Supplementary Table 3.

**Supplementary Table 10:** Calibration slopes (the slope of regressing the true phenotype onto the polygenic score predictor) for the six polygenic prediction methods evaluated in Supplementary Fig. 3 and Supplementary Table 4.

**Supplementary Table 11:** Calibration slopes (the slope of regressing the true phenotype onto the polygenic score predictor) for the six polygenic prediction methods evaluated in Supplementary Fig. 4 and Supplementary Table 5.

**Supplementary Table 12:** Calibration slopes (the slope of regressing the true phenotype onto the polygenic score predictor) for the six polygenic prediction methods evaluated in Supplementary Fig. 5 and Supplementary Table 6.

**Supplementary Table 13:** Information on the genome-wide association summary statistics of the six common complex diseases (breast cancer, coronary artery disease, depression, inflammatory bowel disease, rheumatoid arthritis, and type 2 diabetes mellitus), and six quantitative traits (height, body mass index, high-density lipoproteins, low-density lipoproteins, cholesterol, and triglycerides).

**Supplementary Table 14:** SNP heritability of the six common complex diseases (breast cancer, coronary artery disease, depression, inflammatory bowel disease, rheumatoid arthritis, and type 2 diabetes mellitus), and six quantitative traits (height, body mass index, high-density lipoproteins, low-density lipoproteins, cholesterol, and triglycerides), on the observed scale and the liability scale, estimated using genome-wide association summary statistics and LD score regression.

**Supplementary Table 15:** Numerical values of the prediction accuracy shown in Fig. 2. Polygenic scores were trained with large-scale genome-wide association summary statistics, using the 1000 Genomes Project European sample as an external linkage disequilibrium (LD) reference panel. For each of the curated diseases (breast cancer, coronary artery disease, depression, inflammatory bowel disease, rheumatoid arthritis, and type 2 diabetes mellitus), and quantitative traits (height, body mass index, high-density lipoproteins, low-density lipoproteins, cholesterol, and triglycerides), the Partners HealthCare Biobank sample was repeatedly and randomly split into a validation set comprising 1/3 of the data and a testing set comprising 2/3 of the data. Tuning parameters ($P$-value threshold in P+T, fraction of causal SNPs in LDpred, and global shrinkage parameter in PRS-CS) were selected in the validation data set, and the predictive performance was assessed in the testing set. The mean and standard deviation of each prediction accuracy metric ($R^2$ or Nagelkerke's $R^2$, area under the receiver operating characteristic [ROC] curve, area under the precision-call curve, odds ratio [OR] comparing top 10% of the participants having high polygenic risk with the remaining 90% of the sample) for each polygenic prediction method across 100 random splits are reported.

**Supplementary Table 16:** Numerical values of the prediction accuracy shown in Supplementary Fig. 6. Polygenic scores were trained with large-scale genome-wide association summary statistics, using the Partners HealthCare Biobank data as an in-sample linkage disequilibrium (LD) reference panel. For each of the curated diseases (breast cancer, coronary artery disease, depression, inflammatory bowel disease, rheumatoid arthritis, and type 2 diabetes mellitus), and quantitative traits (height, body mass index, high-density lipoproteins, low-density lipoproteins, cholesterol, and triglycerides), the Partners HealthCare Biobank sample was repeatedly and randomly split into a validation set comprising 1/3 of the data and a testing set comprising 2/3 of the data. Tuning parameters ($P$-value threshold in P+T, fraction of causal SNPs in LDpred, and global shrinkage parameter in PRS-CS) were selected in the validation data set, and the predictive performance was assessed in the testing set. The mean and standard deviation of each prediction accuracy metric ($R^2$ or Nagelkerke's $R^2$, area under the receiver operating characteristic [ROC] curve, area under the precision-call curve, odds ratio [OR] comparing top 10% of the participants having high polygenic risk with the remaining 90% of the sample) for each polygenic prediction method across 100 random splits are reported.
